# Suppression of vitamin D metabolizing enzyme CYP24A1 provides increased sensitivity to chemotherapeutic drugs in breast cancer

**DOI:** 10.1101/2022.10.18.512762

**Authors:** Sakura Kamiya, Nakamori Yuna, Akira Takasawa, Kumi Takasawa, Daisuke Kyuno, Yusuke Ono, Kazufumi Magara, Makoto Osanai

**Affiliations:** Department of Pathology, Sapporo Medical University School of Medicine, Sapporo, Japan; Department of Oral Surgery, Sapporo Medical University School of Medicine, Sapporo, Japan

**Keywords:** Breast cancer, vitamin D, vitamin D metabolizing enzyme CYP24A1, oncogenic property, anticancer drugs, immunohistochemistry

## Abstract

Vitamin D is an essential nutrient for the human body that plays an important role in homeostasis and contributes to cell fate decision including proliferation, differentiation and cell viability. Accumulated epidemiological data suggest a relationship between vitamin D deficiency and carcinogenesis in various organs. However, the underlying mechanism remains to be further clarified. In this study, we showed that vitamin D metabolizing enzyme CYP24A1 promotes the oncogenic property of breast carcinoma cells. In addition, expression of CYP24A1 is elevated in invasive breast carcinoma and decreases the overall survival. Importantly, CYP24A1 suppression significantly enhances cell death sensitivity to two pharmacologically different acting anticancer drugs, cisplatin and gefitinib. The results of our study suggest the possibility of CYP24A1-inhibiting therapy for breast malignancy.

## Introduction

Vitamin D is an essential nutrient for the human body and is not only crucial for regulating calcium metabolism but also plays an important role in homeostasis^1-4^. 1,25-dihydroxyvitamin D (1α,25(OH)_2_D_3_), known as calcitriol, is the active form of vitamin D. Previous evidence has shown that 1α,25(OH)_2_D_3_ is a ligand of nuclear vitamin D receptor that contributes to various processes in the body including cell proliferation, differentiation and viability^5-7^. 1α,25(OH)_2_D_3_ can act protectively against cancer mainly by promoting apoptosis^8^ and the relationships between vitamin D deficiency and various types of cancer such as colorectal and prostate cancer have been explored in several studies^9-11^. In addition, it has been shown that supplementation of vitamin D suppresses carcinogenesis in a variety of organs^12^. However, the underlying mechanism linking tumorigenicity and cellular vitamin D status remains unknown.

The bioavailability of vitamin D is regulated by a coordinated balance between 1α,25(OH)_2_D_3_ biosynthesis and catabolism and determines cellular responses to vitamin D. The vitamin D metabolizing enzyme CYP24A1 (cytochrome P450 family 24 subtype A1) contributes to the inactivation of 1α,25(OH)_2_D_3_ by converting it to rapidly excreted derivatives. This enzymatic activity restricts the access of 1α,25(OH)_2_D_3_ to the transcriptional machinery and limits vitamin D signaling within cells. It is known that CYP24A1 expression is elevated in different types of tumor cells and many of them contain inactive vitamin D metabolites, such as 1α,24,25-(OH)_3_D_3_ and 24 -oxo-1α,25- (OH)2 D_3_^13,14^.

Previous studies showed that CYP24A1 has an oncogenic activity in breast cancer^15^, while the clinical relevance of vitamin D depletion induced by CYP24A1 in breast cancer remains to be clarified. In this study, we investigated the expression of CYP24A1 in surgically resected breast tumor specimens and the effect of CYP24A1 expression on the carcinogenic insults of breast carcinoma cells. The data demonstrated that high expression level of CYP24A1 is related to an increased mortality rate in breast cancer. In addition, suppression of CYP24A1 provides increased sensitivity to chemotherapeutic drugs. Our results suggest CYP24A1 is a potential therapeutic target of breast cancer.

## Materials and methods

### Patients and specimens

Specimens of 136 cases of breast cancer collected by surgical resection during the period from 2011 to 2014 were used in this study. Data were collected from the pathology file of Sapporo Medical University Hospital, Sapporo, Japan. The mean age of the patients was 59.3 years (range, 26-92 years). Histological type was based on the WHO classification of tumors of the breast (4th edition). All of the 136 cases were staged by the Union for International Cancer Control (UICC) stage classification (7th edition) (Table 1). This study was approved by the Institutional Review Board of Sapporo Medical University (IRB study number: 312-230). Written informed consent was obtained from each patient who participated in the investigation. The research was conducted in accordance with the Helsinki Declaration.

**Table 1.**
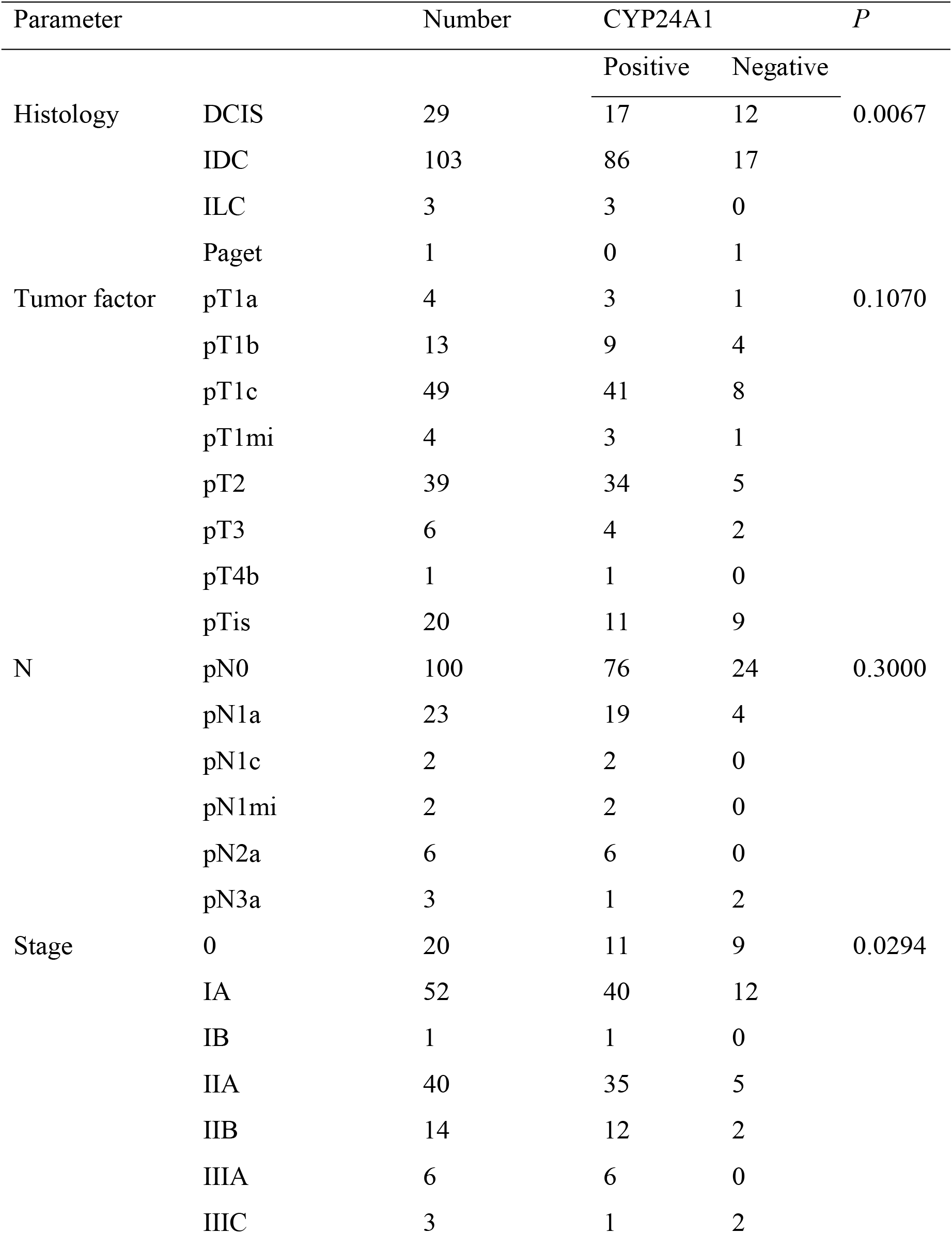
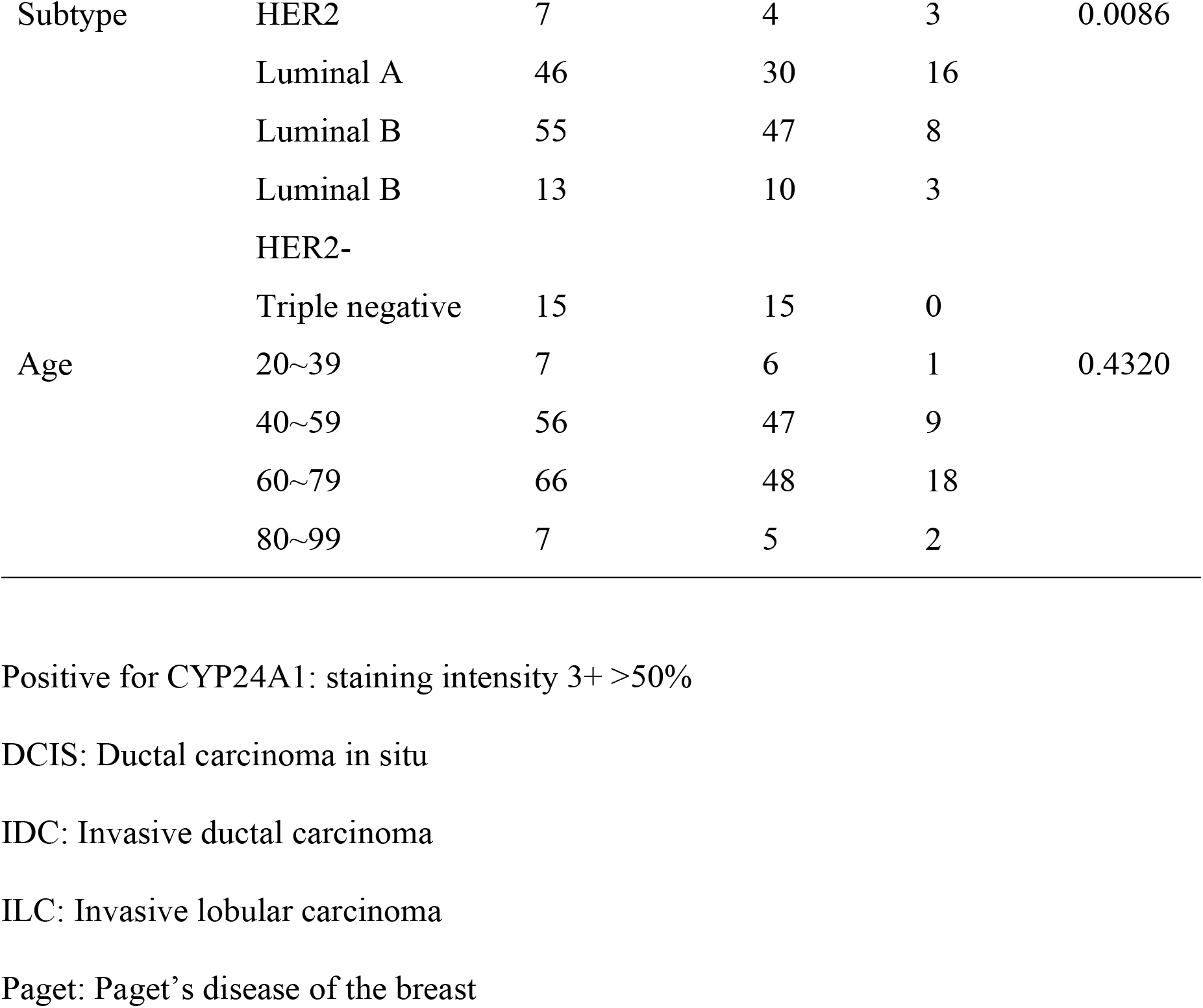
Association between CYP24A1 expression examined by immunohistochemistry and various clinicopathological parameters

### Immunohistochemical staining

Tissue sections were deparaffinized in xylene, rehydrated through a graded series of ethanol and PBS, and incubated in 3% H_2_O_2_ for 10 minutes to block endogenous peroxidase activity. After antigen retrieval by microwave heating (95°C for 30 minutes) in 10 mmol/L Tris/1 mmol/L EDTA buffer, sections were incubated overnight at 4°C with a primary monoclonal antibody against CYP24A1(sc-365700, Santa Cruz Biotechnology Inc, CA, USA. The sections were then incubated with EnVision (Dako, Agilent, Santa Clara, CA, USA) for 30 minutes at room temperature, and 3,3’-diaminobenzidine tetrachloride (Dako) was added as the chromogen. Hematoxylin was used for counterstaining. Analysis of immunohistochemical staining positivity was conducted with consideration of the staining intensity and percentage of positive cells. The intensity scores of staining were set as follows: 3+ strong, 2+ moderate, 1+ weak and 0 negative. The observers were blinded to the clinical data during the evaluation. Consensus was reached by discussing discordant cases.

### Cell culture and transfection

Breast cancer cell line MCF7 cells are estrogen receptor (ER)-positive cells. The cell line was obtained from a local distributor (Summit Pharmaceuticals International, Tokyo, Japan) of the American Type Culture Collection (Manassas, VA, USA). All cells were maintained in Dulbecco’s modified Eagle’s medium (Sigma, St. Louis, MO, USA) supplemented with 10% fetal bovine serum (FBS) (Invitrogen, Carlsbad, CA, USA) and 5% streptomycin (Sigma). The cells were transfected with different types of CYP24A1-specific small-hairpin RNA (shRNA)-expressing lentivirus plasmids (Sigma) using FuGENE6 (Roche, Basel, Switzerland) to generate stable transfectants: CYP24A1 shRNA #1296 (harboring shRNA clone NM_ 000782.2-1296s1c1) and CYP24A1 shRNA #1016 (harboring shRNA clone NM_000782.2-1016s1c1). Transfected clones were selected in 1.5 µg/ml puromycin (Sigma). We selected drug-resistant clones after more than 14 days of selection and screened them for CYP24A1 expression. We obtained the following MCF7 cell transfectants: CYP24A1 shRNA #1 and CYP24A1 shRNA #7.

### Semiquantitative reverse transcription-polymerase chain reaction (RT-PCR) analysis

Total RNA was isolated using the TRIzol reagent (Invitrogen), and subsequent RT-PCR was conducted using the Superscript II Reverse Transcriptase kit (Invitrogen). Samples were incubated at 42°C for 50 min followed by incubation at 70°C for 15 min. Then the cDNA was mixed with appropriate primers and 0.5 U of Taq DNA polymerase (Takara, Shiga, Japan) to amplify the genes of interest. The cycling conditions were set as follows: 30-40 cycles of 30 sec at 96°C, 30 sec at 55°C, and 1 min at 72°C, followed by a final elongation time of 7 min at 72°C. The sequences of the PCR primers used are available upon request. RNA expression was calculated using ImageJ software (National Institutes of Health, Bethesda, MD, USA) with normalization to GADPH expression.

### Immunofluorescent assay

Cells were seeded in 35 mm dishes (Iwaki, Chiba, Japan) containing 15 mm cover glasses coated with rat tail collagen and incubated with 10% FBS. The cells on the cover glass were fixed for 10 minutes with a fixing solution (aceton : ethanol = 1:1) at 20°C. The cells were incubated with a primary monoclonal anti-CYP24A1 antibody (1:100) at 4°C overnight. The cells were then treated with Alexa Fluor 488 (green)-conjugated anti-rabbit IgG (1:200) for one hour at room temperature, and nuclei in the cells were counterstained with 4’,6-diamidino-2-phenylindole (Sigma). The samples were examined by using an epifluorescence microscope (Olympus, Tokyo, Japan).

### Treatment of cells

To evaluate cell viability, cells were seeded in 12-well dishes at a density of 40000 cells per well and cells were counted by manual cell counting with trypan blue dye exclusion in a time and dose-dependent manner using a microscope (Olympus, Tokyo, Japan). To assess cells viability under conditions of different cell stresses, cells were treated with oxidative stress using H_2_O_2_ (0, 25, 50, 75µM) and incubated for 4, 8 and 12 hours. For the assessment of cell proliferation with and without vitamin D (Sigma) and ketoconazole (Sigma), cells were seeded in 12-well dishes at a density of 1000 cells per well. Cells were treated with 1 µM vitamin D and 2 µM ketoconazole and incubated for 48 hours.

For assessment of drug sensitivity, cells were seeded in 96-well plates (5000 cells/well) and treated with cisplatin (Adipogen AG, Liestal, Switzerland) and gefitinib (Cayman Chemical, MI, USA) (concentration ranging from 0 to 200 µmol/L). The viability of cells treated with cisplatin was assessed every 24 hours until 96 hours and the viability of cells treated with gefitinib was assessed after 48 hours. Cell viability was analyzed with the WST8 assay using a CCK-8 (Dojindo Laboratories, Kumamoto, Japan) according to the manufacturer’s instructions. Absorbance at a wavelength of 450 nm was measured by using a Varioskan LUX (Thermo Fisher Scientific).

### Immunohistochemistry of cell blocks

Cells treated with oxidative stress induced by H_2_O_2_ (0, 100, 750 µM) were harvested from culture dishes with a cell lifter and then collected by centrifugation at 300 g for 3 minutes. Following the standard method of paraffin-embedding and sectioning, the collected cells were fixed with 10% agarose gel and fixed in formalin overnight at 4°C. Immunostaining was carried out using antibodies against cleaved caspase-3 (#9664; Cell Signaling Technology, Danvers, Massachusetts, USA) and Ki-67 (MIB-1 clone; BioGenex, California, USA).

### Colony formation

Cells were seeded in 12-well plates at a density of 2500 cells per well. After incubation for 48 hours, the cells were fixed with formalin for 15 minutes and stained with 0.04% crystal violet for 15 minutes at room temperature. Area of colonies were counted using ImageJ.

### Statistical analysis

At least 3 independent experiments were conducted for each analysis and all data are expressed as means ± standard deviations. Fisher’s exact test was used for the analysis of statistical differences. We considered differences with a *P*-value of <0.05 to be statistically significant.

## Results

### CYP24A1 is expressed in primary breast neoplasia

Results of previous studies showed that CYP24A1 is highly expressed in different types of cancer^13,14^. Here, we investigated the correlation between CYP24A1 expression examined by immunohistochemistry and the clinicopathological parameters of breast cancer (Table 1 and Figure 1A). CYP24A1 expression was strongly but only partially observed in normal ductal and acinar cells (data not shown). In non-invasive breast carcinoma, CYP24A1 expression was positive in 58.6% (17/29) of the cases. Consistent with previous findings, the CYP24A1 positive rates were 83.5% (86/103) and 100% (3/3) in invasive ductal carcinoma and invasive lobular carcinoma, respectively (*P*=.0067). Although associations of CYP24A1 expression with primary tumor status (pT) and lymph node involvement (pN) were not found, tumor stage was positively associated with a high expression level of CYP24A1 (*P*=.0294). In addition, intrinsic surrogate subtype was associated with the expression of CYP24A1, being significantly higher in cancer with poor prognosis. Although some are overlapping, the prognosis of breast cancer patients declines in the order of luminal A, luminal B, human epidermal growth factor receptor 2 (known as HER2) overexpressed type, and triple negative type^15-18^. Indeed, a significant increase in the expression of CYP24A1 was noted in hormone negative cancers and luminal B with and without expression of HER2 (*P*=.0086).

**Figure 1.**
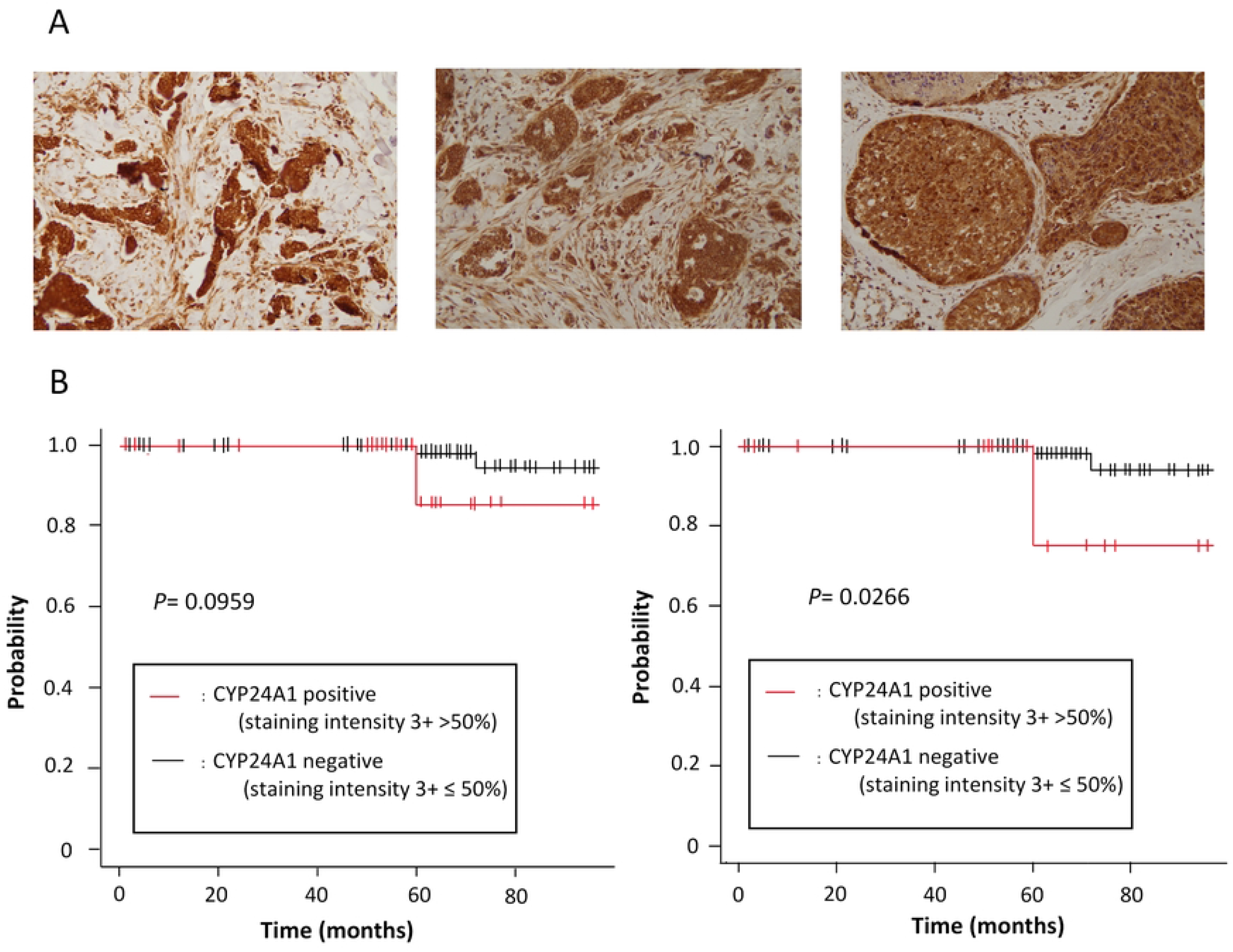
Expression of CYP24A1 in primary breast neoplasia. A. Three representative cases of CYP24A1 positivity (staining intensity 3+ >50%) Original magnification, x200. B. Kaplan-Meier analysis of CYP24A1 positive and negative specimen in whole breast tumor (left panel) and invasive breast cancer (right panel).

The cases were next divided into two groups by the expression of CYP24A1 assessed by staining intensity and area (Figure 1B). Specimens containing >50% area with staining intensity 3+ were defined as positive, and specimens containing ≤50% area with staining intensity 3+ were defined as negative. Kaplan-Meier survival curves showed that the overall survival rate in the CYP24A1 positive group was shorter than the rate in the CYP24A1 negative group when compared in whole specimens. However, a significant association between the CYP24A1 positive group and overall survival rate was observed in invasive breast carcinoma (*P*=.0266) (Figure 1B). These results are consistent with results of previous studies, suggesting a possible oncogenic effect of CYP24A1 in the process of breast neoplasm^19-21^.

### Establishment of CYP24A1 knockdown cells

For the establishment of CYP24A1-suppressed MCF7 cells, cells were transfected with two different sequences of shRNAs against CYP24A1. Finally, we established two cell lines using two different shRNA constructs and denoted the cell lines as CYP24A1 shRNA #1 and CYP24A1 shRNA #7. An immunofluorescent assay showed that the CYP24A1 protein was sufficiently suppressed (Figure 2A). RT-PCR analysis demonstrated that RNA expression was reduced by 75% in CYP24A1 shRNA #7 and that the expression levels of CYP24A1 were efficiently reduced in CYP24A1 shRNA #1 (Figure 2B).

**Figure 2.**
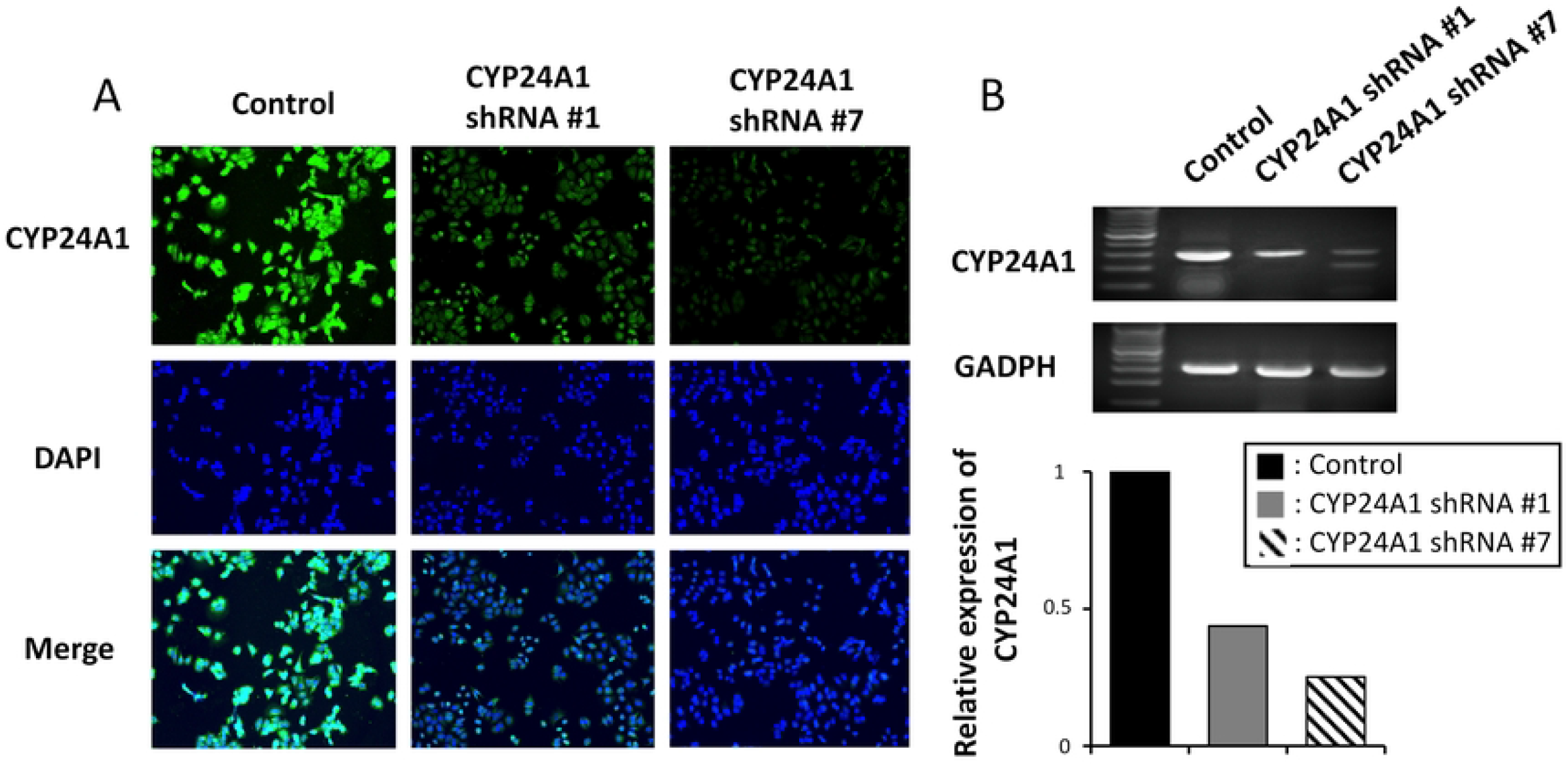
Establishment of CYP24A1 knock down MCF7 cells. A. Immunofluorescent assay of CYP24A1 and DAPI. Original magnification, x100. B. RT-PCR analysis of CYP24A1 (upper panel) and its quantitative analysis (bottom panel) defined as 100% in control cells.

### Effect of CYP24A1 suppression on cell viability

To explore the effect of CYP24A1 knockdown on cell viability, we cultured MCF7 cells for 12 hours with and without oxidative stress using H_2_O_2_ (Figure 3A). In the absence of H_2_O_2_, no difference in cell viability was observed in all cell groups. The cell viability decreased in CYP24A1 shRNA #7 in a dose and time-dependent manner. However, there was no significant difference in CYP24A1 shRNA #1.

**Figure 3.**
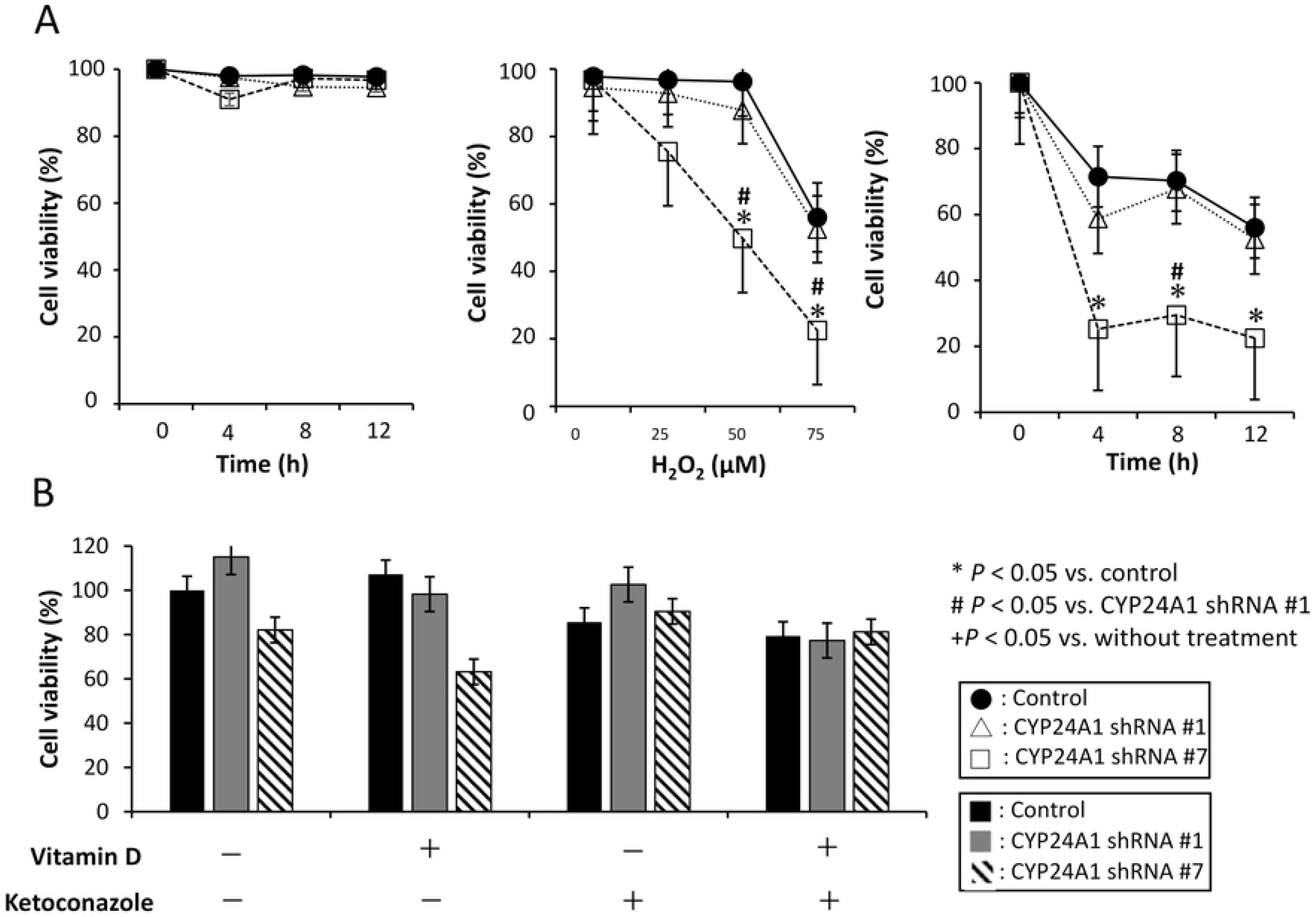
Effect on cell viability with CYP24A1 suppression in MCF7 cells. A. Cell viability without oxidative stress (left panel), with various concentration of oxidative stress such as H_2_O_2_ (25-75 µM) (middle panel), with H_2_O_2_ (75 µM) in a time dependent manner in MCF cells (right panel). B. Quantitative analysis of cell viability with CYP24A1 suppression in MCF7 cells in the presence and absence of vitamin D (1 µM) and ketoconazole (2 µM).

We next treated cells with vitamin D and ketoconazole, a broad-spectrum inhibitor of CYP24A1, and performed a manual cell count after 48 hours. MCF7 cells are ER-positive breast cancer cells which are currently the most commonly used model of breast cancer. Although ER-positive cells are known to express higher levels of vitamin D receptor (VDR) than ER-negative cells^22^, vitamin D only appreciably decreased cell viability in CYP24A1 shRNA #7. There was no difference in cell viability when ketoconazole was added (Figure 3B).

### Effect of CYP24A1 suppression on apoptosis

We established cell blocks from cultured cells to assess cell death sensitivity under the condition of cell stress and counted the number of apoptotic bodies manually (Figure 4A). Apoptosis was significantly increased in CYP24A1 knockdown cells. The number of apoptotic cells increased in CYP24A1-suppressed cells when a moderate level of oxidative stress (100 µM H_2_O_2_) was added. In controls, the number of apoptotic cells only increased with a higher concentration of oxidative stress (750 µM). The data suggest that CYP24A1-suppressed cells have a higher cell death sensitivity to a cell stressor.

**Figure 4.**
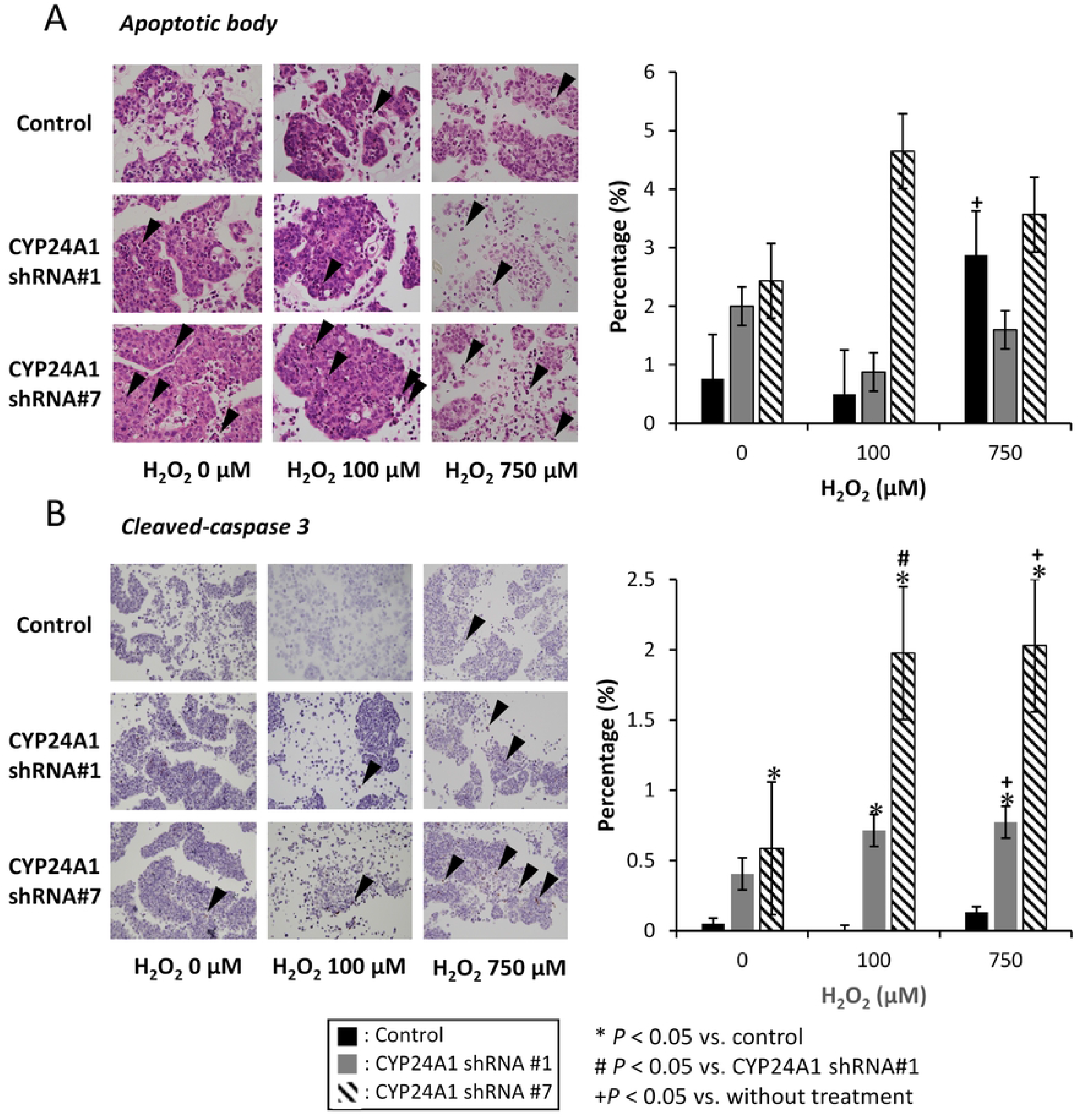
Effect on cell apoptosis with CYP24A1 suppression. Apoptosis was evaluated using cell blocks cultured with different doses of H2O2 (100, 750 µM). A. Manual cell counting of apoptotic bodies. B. Immunohistochemistry of cleaved caspase-3. Representative images (left panels), original magnification x200 (A) and x100 (B). Quantitative analysis (right panel).

We next carried out immunohistochemistry using an antibody against cleaved caspase-3 to evaluate the effects of altered CYP24A1 expression on apoptosis (Figure 4B). Cleaved caspase-3-positive cells were significantly increased in CYP24A1 knockdown cells treated with H_2_O_2_ in a dose-dependent manner.

### Effect of CYP24A1 suppression on colony forming efficacy

To clarify the effect of CYP24A1 knockdown on two dimensional tumorigenicity with and without vitamin D and ketoconazole, we conducted a colony formation assay (Figure 5). Compared to the colony forming ability without treatment, the ability was suppressed in both CYP24A1 shRNA #1 and shRNA #7. We found that the colony formation efficacy was suppressed more in shRNA #7. In the presence of vitamin D, the area of the colonies was decreased in all cell groups. The results suggest that cellular vitamin D status determines colony formation efficacy. However, ketoconazole did not affect colony formation.

**Figure 5.**
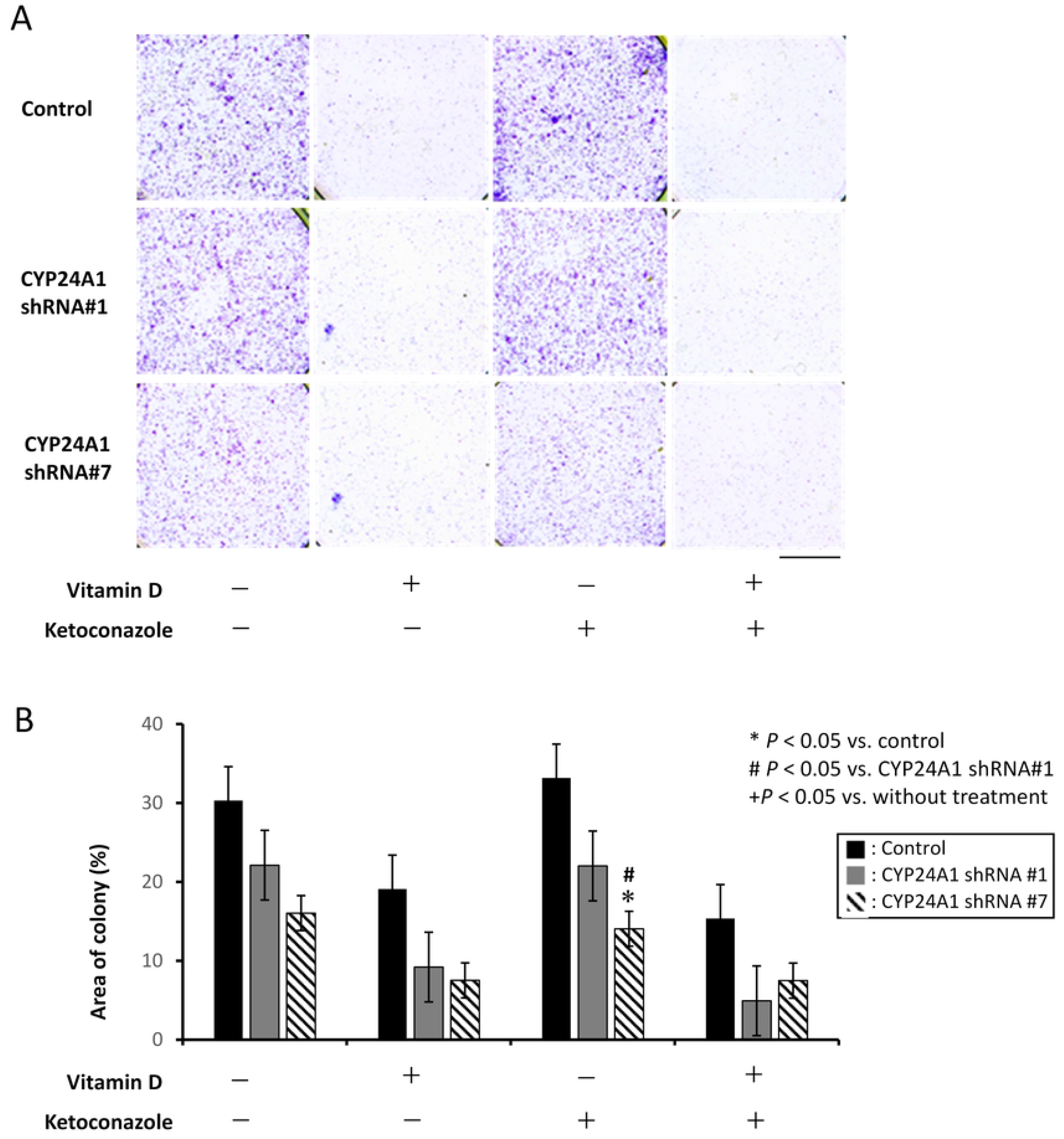
Effect on colony forming efficacy with CYP24A1 suppression. Colony forming efficacy (A) and its quantitative analysis (B) with and without vitamin D (1 µM) and ketoconazole (2 µM).

### Effect of CYP24A1 suppression on cell death sensitivity to anticancer drugs

To investigate the cell sensitivity to anticancer drugs with different pharmacological mechanism (cisplatin and gefitinib), we cultured cells with each drug and analyzed the cell viability (Figure 6). Interestingly, reduced expression of CYP24A1 significantly enhanced cell death sensitivity to both to cisplatin and gefitinib.

**Figure 6.**
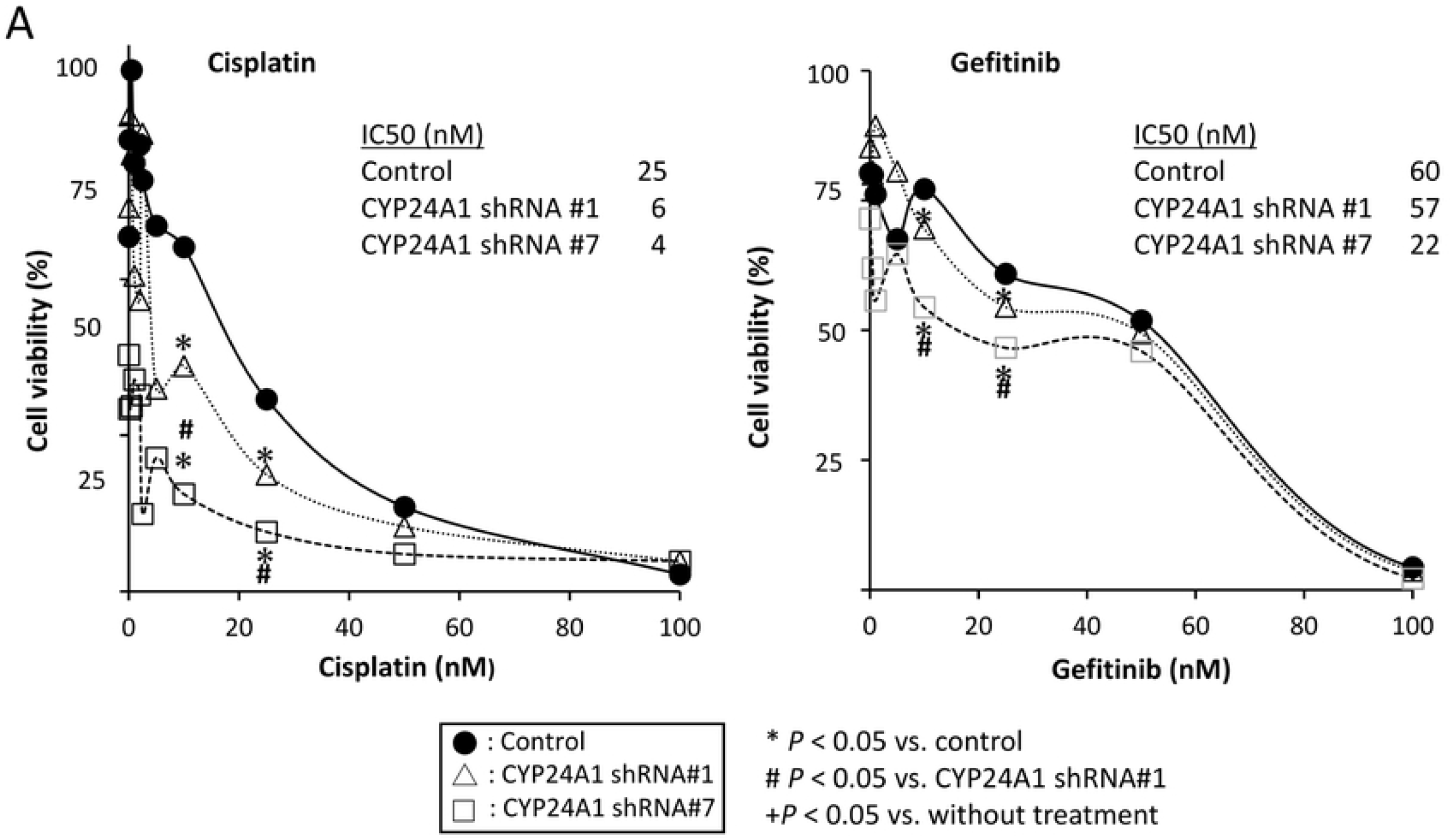
Effect on cell death sensitivity to anticancer drugs with CYP24A1 suppression. MCF7 cells were treated with cisplatin (left panel) and gefitinib (right panel) in a dose dependent manner. We calculated half maximal (50%) inhibitory concentration (IC50).

## Discussion

In this study, we demonstrated for the first time that increased expression of CYP24A1 leads to a decrease in the overall survival in patients with invasive primary breast carcinoma. In addition, suppression of CYP24A1 inhibits oncogenic activity of breast carcinoma cells and enhances cell sensitivity to anticancer drugs with different pharmacologic activities. Our results suggest that CYP24A1 is a possible therapeutic target for breast malignancy.

According to the expression pattern of hormone receptors examined by immunohistochemistry, including estrogen receptor (ER), progesterone receptor (PR) and HER2, breast tumors can be classified in three major immunohistochemically surrogate intrinsic subtypes. The prognosis and treatment response differ among the breast cancer subtypes, triple negative cancer having the worst prognosis and Luminal A the best prognosis^15,16,18^. Previous data showed that ER+ breast cancer cell lines are more sensitive to the effects of calcitriol. In our study, the expression level of CYP24A1 was higher in the intrinsic subtypes with a poor prognosis, particularly in triple negative cancer (Table 1), suggesting that CYP24A1 is a possible marker of prognosis in breast cancer. Supporting the hypothesis, patients with strong expression of CYP24A1 in invasive ductal carcinoma had a low overall survival rate compared with that in patients with those with only moderate or no expression of CYP24A1 (Figure 1).

Different studies have showen that 1α,25(OH)_2_D_3_ has activity for suppressing carcinogenesis and that CYP24A1-mediated intracellular vitamin D metabolism promotes oncogenic behavior^19-21,23^. Our data demonstrated that supplementation of vitamin D decreased cell viability and colony forming efficacy. However, the results were not statistically significant, suggesting that CYP24A1 has an as-yet-unrecognized activity independent of vitamin D metabolism. Furthermore, the addition of ketoconazole, which is a broad-spectrum inhibitor of cytochrome P450^24,25^, did not affect the viability and colony forming efficacy of MCF7 cells, although suppression of CYP24A1 itself significantly decreased these effects (Figure 5). One possible explanation is that a CYP24A1-specific inhibitor is required to effectively inhibit the tumorigenicity of breast cancer.

Our data revealed that CYP24A1 suppression in breast cancer cells increased cell sensitivity to two anticancer drugs with different pharmacological mechanisms. The first anticancer drug we used was cisplatin, a chemotherapeutic drug that induces cell apoptosis in cancer cells by crosslinking with the purine bases on the DNA and disrupting its repair mechanism^26^. The second anticancer drug was gefitinib, which is a tyrosine-kinase inhibitor used for a variety of cancers including HER2-positive breast cancer^27^. Luminal breast cancers are treated with hormone therapy and several studies have demonstrated that ER-positive tumors respond poorly to conventional chemotherapy^15,16^. Our results indicate that CYP24A1 enhances cell death activity to differently acting cell death inducers. Therefore, we propose that a strategy to inhibit CYP24A1 activity is a possible therapeutic entity for breast malignancy.

A limitation of this study is that we only used luminal intrinsic subtype breast cancer cells, since ER-positive cells have been shown to express higher levels of VDR than ER-negative cells^22^ and vitamin D deficiency is associated with poor outcomes in luminal-type breast cancer patients^28^. Results of previous studies also showed that dietary intake of vitamin D reduces the risk of ER-positive breast cancer^29-32^. Notably, our study showed that the expression of CYP24A1 was also significantly higher in triple negative cancer, and further studies using different cell lines with various expression levels of ER, PR and HER2 are needed.

Our study showed that suppression of CYP24A1 in invasive breast cancer leads to an increase in the overall survival rate of patients with breast carcinoma. Supporting this result, we found that suppression of CYP24A1 in vivo decreases tumorigenicity of breast carcinoma cells and increases cell sensitivity to differently working anticancer drugs. In conclusion, the results of our study suggest that CYP24A1 is a possible therapeutic target for breast malignancy.

## Acknowledgments

The works in this study were supported in part by grants from the Grants-in-Aid for Scientific Research Program from the Japan Society for the Promotion of Science (JSPS KAKEHHI) Grant Numbers: 22K06983, 21K08715, 20K07409 and 20K16196

## Conflict of interest

The authors declare that they have no conflicts of interests.

## Notes

### Competing Interest Statement

The authors have declared no competing interest.

